# DNA Binding by the *Escherichia coli* MinC–MinD Complex Requires the MinC Interdomain Linker

**DOI:** 10.1101/2025.02.06.636808

**Authors:** Navaneethan Palanisamy, Linda Klauss, Maïween Lefebvre, Jungsheng Liang, Götz Hofhaus, Johan Zeelen, Maximilian Urban, Mehmet Ali Öztürk, Sabine Merker, Marcin Luzarowski, Thomas Ruppert, Barbara Di Ventura

**Author notes:** Correspondence to: Barbara Di Ventura. current address: Chester Medical School, University of Chester, Chester, UK. current address: Gyros Protein Technologies AB, Uppsala, Sweden. These authors contributed equally (Navaneethan Palanisamy, Linda Klauss).

## Abstract

The *Escherichia coli* Min system, composed of the proteins MinC, MinD, and MinE, is crucial for directing septum formation at mid-cell. MinD has also been shown to bind DNA in a non-sequence-specific manner and to tether plasmid DNA to liposomes. A role for the Min system in chromosome segregation has been proposed, whereby repeated, transient membrane-tethering events mediated by MinD could help prevent backward movements of the duplicated chromosomes once entropic forces weaken. However, the contribution of the other Min proteins to DNA binding has remained unclear.

Here, we demonstrate that DNA binding is strongly enhanced when MinD and MinC form a complex, while MinE and FtsZ (at high concentrations) interfere with this binding. We reveal a critical role for the linker connecting MinC’s N- and C-terminal domains in both DNA binding and inhibition of FtsZ polymerization. *E. coli* strains expressing MinC linker variants from the native locus, in combination with the alpha subunit of HU tagged with GFP, display chromosomes that appear more spread throughout the cell. This phenotype could reflect reduced compaction, chromosome segregation defects, or a combination of both.

These findings suggest a previously unrecognized role for the MinC linker in DNA binding and provide new insights into a potential physiological role of MinCD-DNA bnding in chromosome organization and/or dynamics.

## Introduction

*E. coli* cells divide by synthesis of new cell wall and invagination of the cytoplasmic membrane at mid-cell, which gradually constricts to form two daughter cells^1^. The divisome at mid-cell is assembled around a structure called the Z-ring, which is made of polymers of FtsZ, a conserved GTPase that is structurally similar to eukaryotic tubulin^2, 3^. In the absence of FtsZ’s spatial regulators, the Z-ring would form anywhere in the cell. To inhibit Z-ring formation at the poles and over the nucleoid two machineries are at place in *E. coli*: the Min system and the nucleoid occlusion^4^. ZapA and ZapB have also been shown to localize at mid-cell in an FtsZ-independent manner, and to provide, together with MatP, additional positional cues for Z-ring formation^5–7^. The nucleoid occlusion mechanism is put in action by the protein SlmA, which binds DNA at specific sites scattered along the chromosome with the exception of the terminus-containing region and inhibits FtsZ polymerization there^8, 9^. The Min system comprises three proteins: MinC, MinD and MinE. It owes its name from the observation that *E. coli* strains having mutations in or deletion of the operon encoding these proteins –the *minB* operon– divide asymmetrically giving rise to miniaturized cells deprived of chromosomal DNA^10–12^. These round, chromosome-less cells have been denominated minicells and the proteins forming the machinery that ensures symmetric cell division and absence of minicells were called Min proteins, and together the Min system.

MinC is a potent inhibitor of FtsZ polymerization. While it is able to do so on its own, a 25-50 times higher cellular concentration of MinC is needed to antagonize FtsZ efficiently in the absence of MinD^13^. Indeed, MinC is *per se* a cytoplasmic protein^14, 15^ and, if expressed in the absence of MinD and MinE, it is simply diffused throughout the cell, thus requiring high concentrations to counteract FtsZ, which is instead tethered to the cytoplasmic membrane by its binding partners FtsA and ZapA^16–18^. Thanks to its association with MinD, an ATPase that associates with the cytoplasmic membrane through its C-terminal amphipathic helix (also called a membrane-targeting sequence or MTS)^19–23^, MinC gets in close proximity to FtsZ and can potently destabilize its polymers even at endogenous expression levels. MinC is composed of two functional domains: an N-terminal domain (MinC^N^) and a C-terminal domain (MinC^C^) connected by a linker allowing flexibility in the orientation of MinC^N24, 25^. The boundaries of these domains are not universally defined. Some reports define MinC^N^ as the region of the protein comprising amino acids 1-120, while MinC^C^ as comprising amino acids 127-231^26^; others state MinC^N^ to comprise amino acids 1-115 of MinC and MinC^C^ to include amino acids 116-231^27^. So far, nothing is known about the role of this linker except that it imparts flexibility between the two domains of MinC. Mutations in this region impairing MinC’s functionality have never been identified. MinC^C^ contains a dimerization domain and interacts directly with MinD^28^. When in complex with MinD, MinC^C^ interacts with the C-terminal tail of FtsZ, which is also involved in the interaction with FtsA and ZapA^29^, thus competing with these proteins and finally destabilizing and disrupting the Z-ring^29, 30^. MinC^N^ is also dimeric due to a so-called domain swap, whereby the first beta-strand of each monomer belongs to a beta-sheet located on the other monomer^31^. MinC^N^ is a stronger FtsZ antagonist than MinC^C^. It attacks and shortens FtsZ polymers without affecting FtsZ’s GTPase activity^27^. Taken together, it can be concluded that MinC binds to FtsZ and destabilizes its polymers with both its N- and C-terminal domains^24, 32^.

MinD forms ATP-dependent dimers^33^. Although ADP-bound monomeric MinD can associate with the cytoplasmic membrane via its MTS, this interaction is weak. When in dimeric form, the interaction becomes stable^19^. In the absence of MinE or when expressed at much higher levels than MinE, MinD decorates the whole cytoplasmic membrane^14^. MinE is recruited to the membrane by membrane-bound MinD, which triggers a conformational change in MinE that exposes a previously cryptic MTS, which allows MinE to also directly associate with the membrane^34^. Subsequently, MinE stimulates the ATPase activity of MinD, which leads to the dissociation of the MinD dimer and the release of MinD from the membrane^35, 36^. In the cytoplasm, ADP is exchanged with ATP, and ATP-bound MinD rebinds to the membrane at locations where MinE concentration is the lowest, that is, the opposite pole. This reaction-diffusion mechanism leads to oscillations of MinD/MinE from one cell pole to the other^14, 15^. MinE and MinC compete for the same binding surface on MinD^37^. As a matter of fact, MinE displaces MinC from MinD before activating MinD’s ATPase activity^33, 38–40^.

Beyond its role in Z-ring positioning, the Min system has been implicated in chromosome segregation^41^. MinD was shown to bind DNA *in vitro* in a non-sequence-specific manner^41^. A mutant MinD capable of DNA binding, but unable to bind the membrane, was shown to co-localize with the nucleoid in living *E. coli* cells^41^. MinD was found to bind the DNA and liposomes at the same time^41^. Later, the DNA-binding protein Noc, the actuator of the nucleoid occlusion mechanism in *Bacillus subtilis*, was also shown to bind simultaneously to the membrane and the DNA, suggesting that such “dual binders” might be more widespread than previously thought^42^. MinD is part of the ParA-MinD family of ATPases^23, 43^. Members of this family are known to bind DNA to perform either plasmid or chromosome segregation in various organisms^44^. MinD has been seen for a long time as the exception, because it was believed to bind the membrane exclusively and not DNA, and to function solely in mid-cell determination. Interestingly, *E. coli* does not possess a chromosomal Par system and chromosome segregation is believed to occur spontaneously as a consequence of entropic forces^45, 46^. Nonetheless, proteins that bind the nucleoid could support such an entropy-driven segregation mechanism by regulating physical properties of the chromosomes^46^.

As a matter of fact, in the mechanism proposed by Di Ventura and colleagues^41^, the Min system is said to support chromosome segregation after entropic forces become weaker due to increased separation of the duplicated chromosomes. Thanks to repeated, transient membrane-tethering events mediated by MinD, chromosomes would be prevented from randomly moving backward towards mid-cell^41^.

So far, only MinD-DNA interaction was directly studied by various biochemical assays and fluorescence microscopy in living *E. coli* cells^41^. Here we investigate the interaction of MinC and MinE with DNA *in vitro*, alone or when together with the other Min proteins. We show that DNA binding is enhanced when MinC and MinD form a complex, and that MinE and FtsZ (only at high concentrations) interfere with the binding. Using negative-stain electron microscopy, we show that MinC and MinD form sheets in presence of DNA, with the width of the sheets proportional to the length of the DNA, and that gold beads coupled to double-stranded DNA align along MinCD co-polymers. We find that the linker connecting MinC^N^ and MinC^C^ is essential for MinCD-DNA binding, which is abolished *in vitro* by even small perturbations to it. The linker is also important for MinC’s function towards FtsZ. Using the CRISPR/Cas9 and Lambda Red recombineering^47^, we construct MG1655 and MG1655-HUαGFP (Fishov)^48^ strains expressing MinC linker mutants from the endogenous locus. While in MG1655 there is no clear phenotype, some of the Fishov mutant strains display chromosomes that appear more spread throughout the cell. This phenotype could reflect reduced compaction, chromosome segregation defects, or a combination of both. These data suggest that MinCD-DNA binding has a physiological role and reveal an unexpected role for the MinC linker in the binding to DNA and FtsZ.

## Results

### The MinCD complex binds DNA *in vitro*

To test the ability of MinC, alone and with MinD, to bind DNA, we cloned the *minC* and *minD* genes in pET28a and purified the 6xHis-T7-tagged proteins from *E. coli* Rosetta cells using Nickel-based affinity chromatography. We then performed electrophoretic mobility shift assays (EMSAs) using a fluorescently labeled 30 bp DNA fragment. While MinC and MinD alone bound DNA extremely weakly (Fig. S1a,b), the MinCD complex bound DNA strongly, in a concentration-dependent manner in the presence of ATP (Fig. S1b). When we used a longer DNA fragment (120 bp), we found that His-MinC at 2 μM could also shift all the DNA probe (Fig. S1c), albeit in this case the nucleoprotein complex was found mostly in the well and not into the gel, as seen for the sample with MinC and MinD (Fig. S1d).

These EMSAs were performed with the His-tagged proteins. To test if the His tag could influence the binding, we removed it from both proteins using thrombin and performed the EMSAs again (note that the T7 tag is still present at the N-terminus of the proteins). We found that, indeed, both proteins had reduced DNA binding in the absence of the His tag, yet the binding was still strong when using both proteins together, especially when using a longer DNA probe (Fig. S1e and Fig. 1a). Importantly, this strong binding was observed only when co-incubating MinC and MinD: when MinD was co-incubated with MinE, there was only a mild band visible at high MinE concentrations (Fig. 1b). We decided to perform all further experiments with purified proteins from which the His tag was removed.

**Fig 1.**
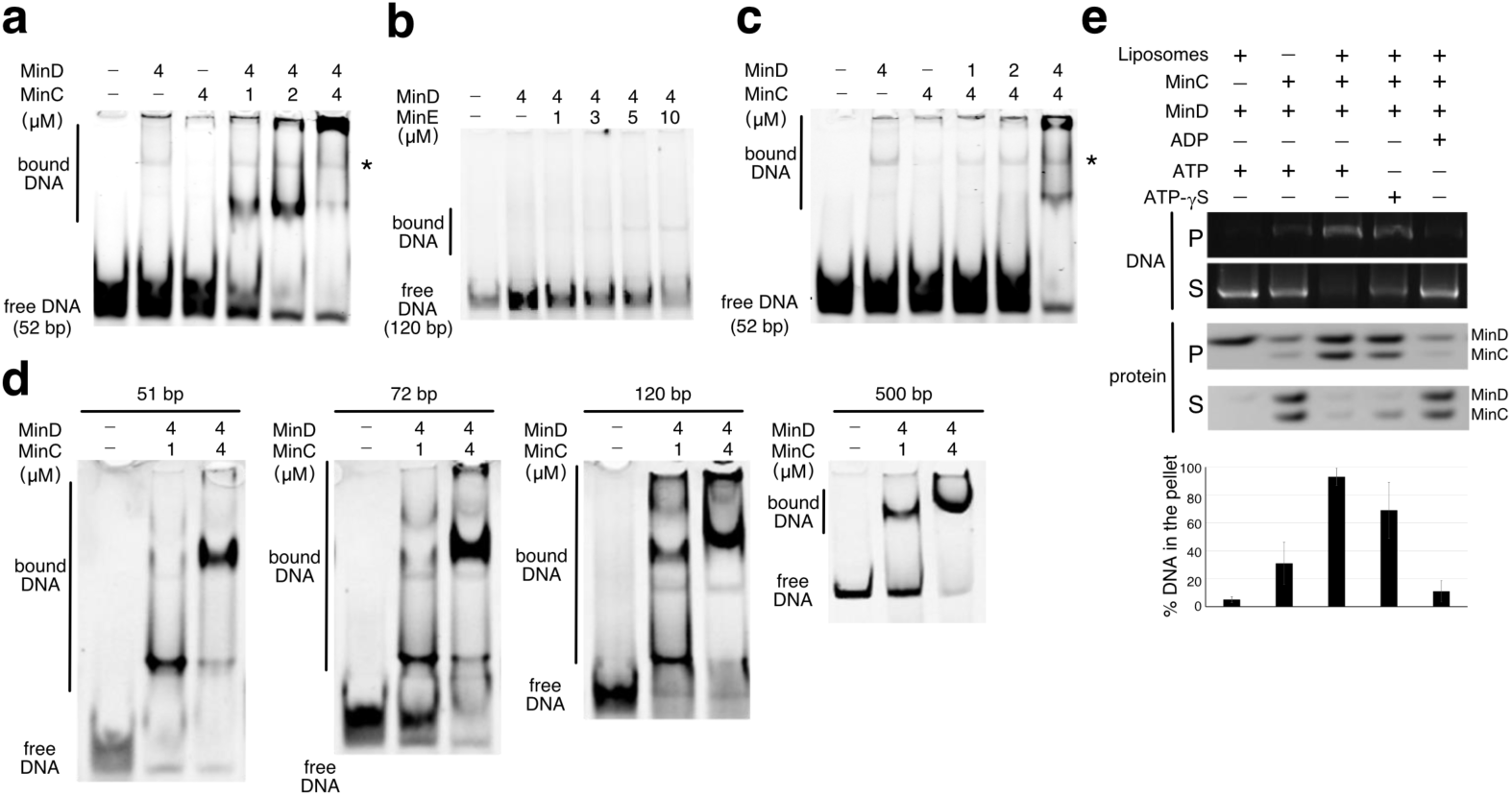
The MinCD complex binds DNA *in vitro*. **a-d,** Electrophoretic mobility shift assays (EMSAs) performed with the indicated proteins at the indicated concentrations, and HEX-labelled DNA (50 nM) of the indicated length. All reactions were performed in the presence of 1 mM ATP. **a**,**b**, Asterisk indicates the height at which a band is visible, which is observed when incubating the DNA with MinD or MinC alone and does not represent binding by the MinCD complex. **e**, Liposome co-sedimentation assay. Upper panel, gel red-stained agarose gel used to visualize the DNA. Middle panel, Coomassie-stained polyacrylamide gel used to visualize the proteins. Lower panel, bar graph showing the quantification of the % of the DNA found in the pellet. Data represent mean ± S.E.M. of three independent experiments. S, supernatant. P, pellet.

Titrating the proteins one at a time with a 52 bp DNA probe we saw that, when having MinD at 4 μM, 1 μM MinC was sufficient to shift the vast majority of the DNA probe (Fig. 1a), while, when having MinC at 4 μM, 4 μM MinD were necessary to observe a similar strong binding (Fig. 1c). When we used a longer DNA probe, we observed the same pattern, although in this case, 2 μM MinD were sufficient to shift all the DNA probe (Fig. S1f,g). Taken together, these data indicate that both, MinC and MinD have rather weak affinity for DNA when alone, but this affinity increases several folds when the proteins form a complex. The concentration-dependent binding of the MinCD complex with DNA was observed with probes of various lengths (Fig. 1d).

Next, we wanted to know if the MinCD complex can tether DNA to liposomes. Therefore, we performed a liposome co-sedimentation assay in the presence of plasmid DNA. We found that the plasmid DNA co-sedimented with the liposomes when MinC and MinD were mixed together with ATP (Fig. 1e, lane 3, upper panel). The MinCD-DNA complex was partially found in the pellet even in the absence of the liposomes (Fig. 1e lane 2, upper panel). In the presence of ADP, there was only a very weak signal for the DNA in the pellet (Fig. 1e, lane 5, upper panel). While this could be due to binding to MinD alone, considering that there was hardly any DNA in the pellet of the sample with only MinD (Fig. 1e, lane 1, upper panel), we speculate that the weak signal is rather due to some residual MinCD complex in the pellet of the sample with ADP (Fig. 1e, lane 5, middle panel). When the non-hydrolysable ATP-analogue ATP-γS was used instead of ATP, we found a decreased amount of DNA in the pellet (Fig. 1e, lane 4, upper panel). Looking at the amount of MinC and MinD in the pellet/supernatant (Fig. 1e, lane 4, middle panel), we conclude that ATP-γS affects the binding between MinC and MinD more than the MinD-liposome interaction, leading to reduced DNA binding. These results indicate that the MinCD complex is capable of binding DNA while being associated with liposomes.

### In the presence of DNA, the MinCD co-polymers assemble *in vitro* into sheets whose width is proportional to the length of the DNA fragment

The results of the liposome co-sedimentation assay, which showed that a fraction of the MinCD-DNA complex goes to the pellet even in the absence of liposomes, suggest that the proteins form a high-molecular weight complex with the DNA. This is also evident in the EMSAs, where the signal is often found in or very close to the well. To more directly observe these structures, we performed negative-stain transmission electron microscopy. Upon incubation of MinC, MinD, ATP and DNA, we observed the formation of sheets (Fig. 2a). We believe these sheets are formed by a number of the previously reported MinCD co-polymers^49,50^. Using DNA fragments of different lengths as well as plasmid DNA, we found that the width of the sheets is proportional to the length of the DNA fragment used in the experiment (Fig. 2b). In the absence of added DNA, sheets are still visible, but they are found more sporadically (Fig. S2).

**Fig 2.**
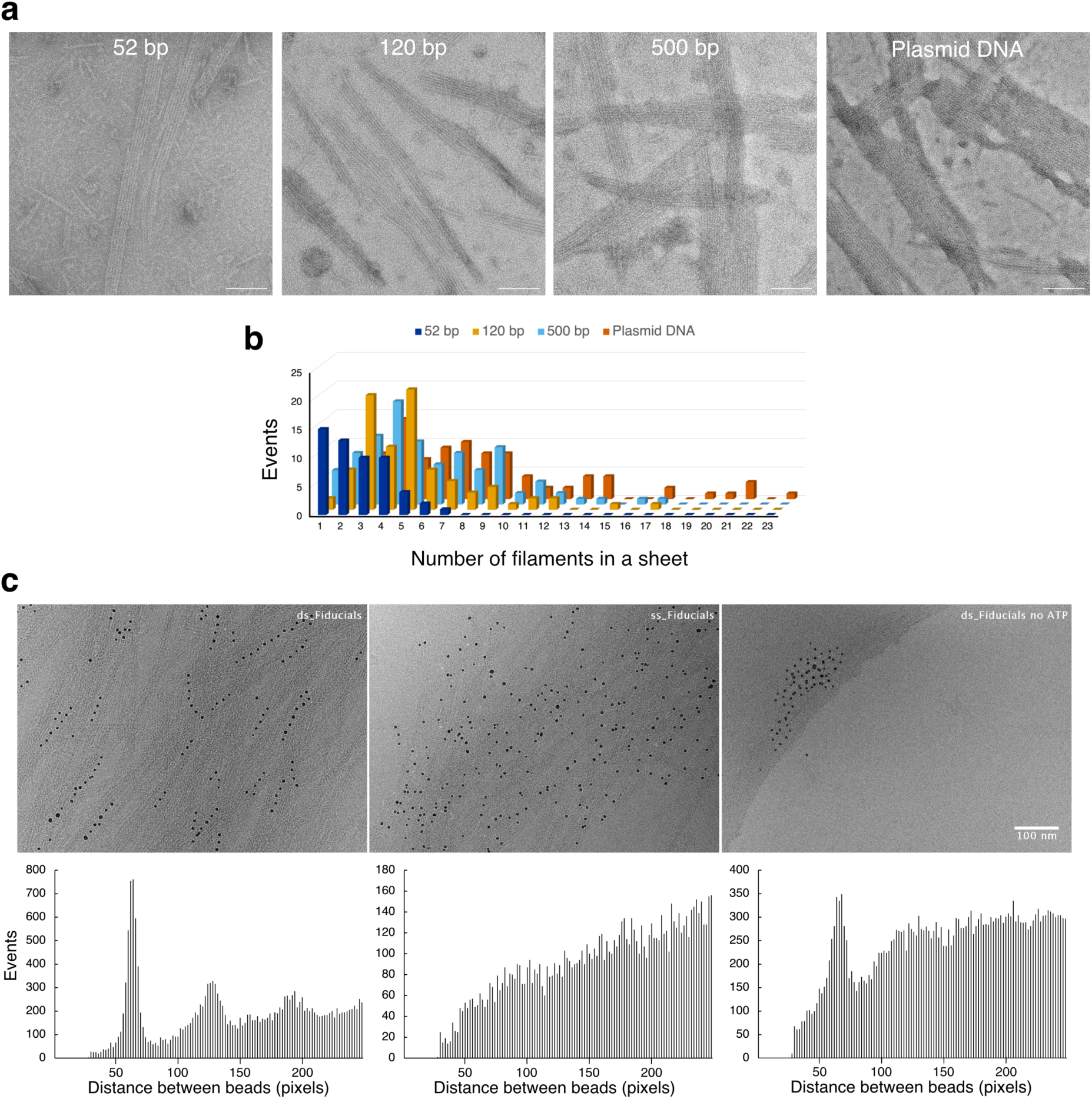
The MinCD co-polymers assemble into sheets in the presence of DNA *in vitro*. **a,** Negative-stain electron-micrographs of samples containing MinC (4 μM), MinD (4 μM), 1 mM ATP, 10 mM MgCl2 and DNA (50 nM) of the indicated length or plasmid DNA. Scale bar, 100 nm. **b**, Bar graph showing the distribution of the number of filaments per sheet for the indicated DNA fragments. **c,** Negative-stain electron-micrographs of samples containing MinC (4 μM), MinD (4 μM), 1 mM ATP and either ds-fiducials (ds DNA) or ss-fiducials (ss DNA). Pixel size, 2.8 Å.

In the experiment described above, the DNA was not directly visible. To visualize the DNA, we used 5 nm gold beads coated with poly-dT DNA via a sulfur-gold chemistry. These single-stranded fiducials (ss fiducials) were annealed with an excess of poly-dA to form double-stranded fiducials (ds fiducials). The MinCD co-polymers were formed by mixing purified MinD and MinC proteins in the presence of ATP. After the addition of the fiducials, the samples were plunge-frozen and observed in the electron microscope. We found that the ds fiducials aligned along the co-polymeric filaments in a linear way (Fig. 2c, left panel) in contrast to the random binding pattern of the ss fiducials (Fig. 2c, middle panel). This regular pattern required the presence of the MinCD filaments (Fig. 2c, right panel). The ss fiducials sometimes formed clusters, which could be reduced in number, but not eliminated, by one additional heating/annealing cycle. This treatment did not affect the regular line pattern seen for the ds fiducials.

A statistical analysis of all inter-fiducial distances up to 250 pixels confirmed the visual observations. The lines of ds fiducials appear to consist of beads at defined distances, and clear peaks corresponding to multiples of this distance (manifolds) are evident in the histogram (Fig. 2c, lower left panel). In contrast, the distances between ss fiducials are random (Fig. 2c, lower middle panel). While the overall binding of ss and ds fiducials to the filaments is comparable, the ss fiducials tend to be more dispersed, resulting in fewer short-range interactions and the disappearance of the peak at the primary distance. The few clusters of ss fiducials observed in the absence of filaments also exhibit a peak at a short distance (18.5 nm), suggesting a potential intrinsic tendency for these fiducials to form small aggregates. These results, together with our EMSA data, support the conclusion that MinC and MinD bind to double-stranded DNA *in vitro*, forming ordered structures.

### MinE and FtsZ interfere with DNA binding by the MinCD complex

Next we investigated the effect of MinE on DNA binding by the MinCD complex using the liposome co-sedimentation assay. We found that MinE interferes with DNA binding by the MinCD complex in a concentration-dependent manner (Fig. 3a). Since MinE competes with MinC for the same binding surface on MinD and dislodges MinC from MinD, it is not surprising that it leads to the dissociation of the MinC-MinD-DNA complex, which requires MinC and MinD to be in a complex.

**Fig 3.**
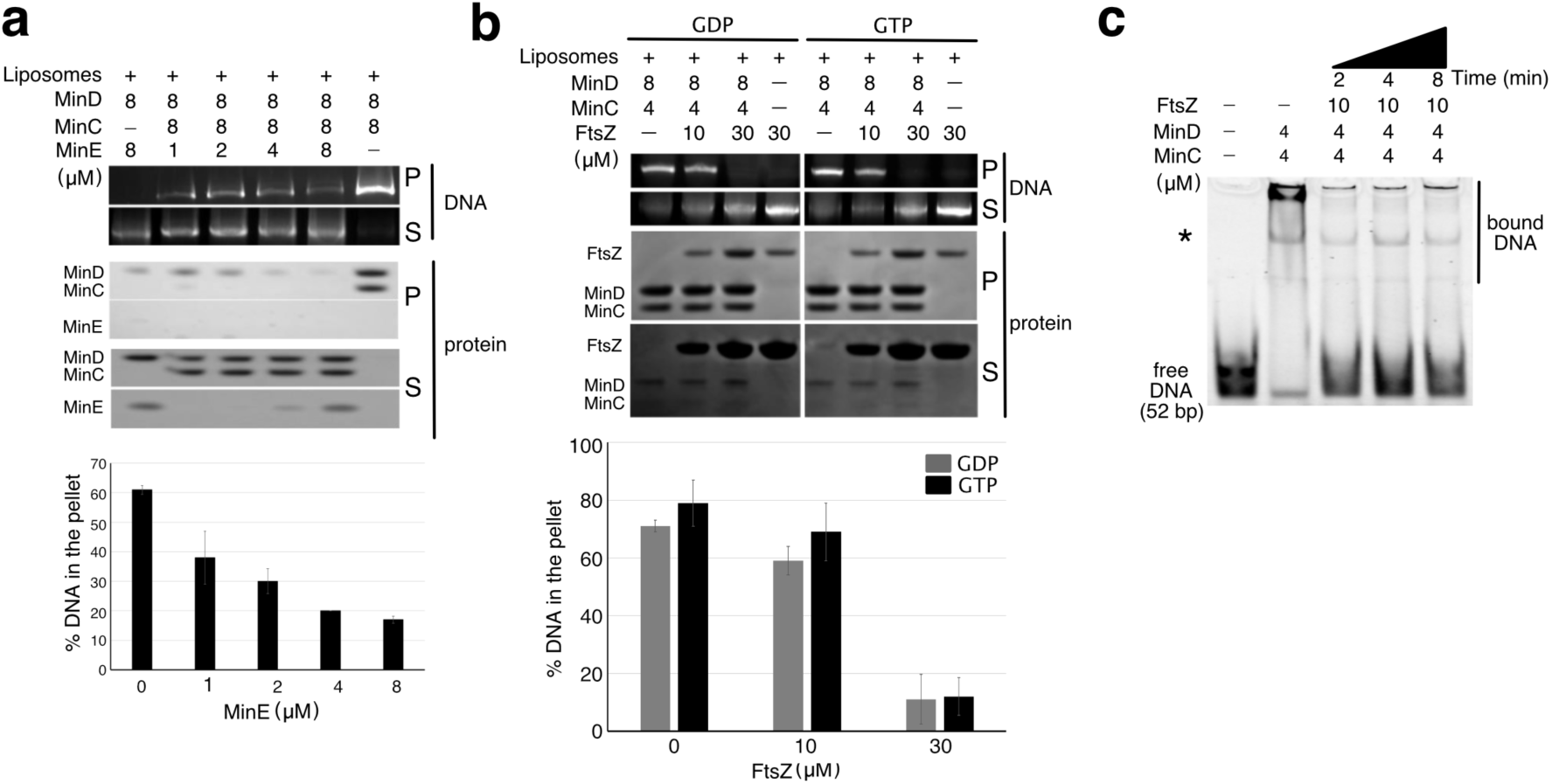
MinE and FtsZ interfere with DNA binding by the MinCD complex. **a-b,** Liposome co-sedimentation assays. Upper panel, Gel-red-stained agarose gel used to visualize the DNA. Middle panel, Coomassie-stained polyacrylamide gel used to visualize the proteins. Lower panel, bar graph showing the quantification of the % of the DNA found in the pellet. Data represent mean ± S.E.M. of three (**a**) and two (**b**) independent experiments. In (**a**), the sample corresponding to 4 μM MinE was measured only once. S, supernatant. P, pellet. **b**, GDP and GTP were added at a final concentration of 1 mM. **c,** Electrophoretic mobility shift assays (EMSAs) performed with the indicated proteins at the indicated concentrations, and HEX-labelled DNA of the indicated length. The time indicates the point at which FtsZ was added to the reaction. Total reaction time, 10 minutes. The asterisk indicates the height at which a band is visible, which is observed when incubating the DNA with MinD or MinC alone and does not represent binding by the MinCD complex.

We also tested if FtsZ would interfere or enhance DNA binding by MinCD. To this aim, we cloned the *ftsZ* gene into pET28a, purified the protein from *E. coli* Rosetta cells using Nickel-based affinity chromatography and removed the His tag. We then performed liposome co-sedimentation assay in the presence of plasmid DNA. FtsZ was found in the pellet either when alone or when co-incubated with MinC and MinD, albeit the amount of protein in the pellet was higher for the sample with MinCD (Fig. 3b, compare lanes 3 and 4, middle panel). There was no discernible difference in the results obtained with GDP and GTP (Fig. 3b). We speculate this is due to the intrinsically high GTPase activity of FtsZ^51^, which leads to the accumulation of GDP in the sample that contained initially only GTP. Note that, in this assay, it is not possible to discern polymers that form in the presence of GTP from shorter oligomers and/or condensates, which form in the presence of GDP^52^. Upon incubation with 10 μM FtsZ, the protein ratio MinC/FtsZ in the lipid-associated pellet was ± 2:1 and there was a small decrease of DNA in the pellet (Fig. 3b, lower panel). Increasing the concentration of FtsZ to 30 μM resulted in an equal amount of MinC and FtsZ in the pellet and highly reduced plasmid DNA in the pellet (Fig. 3b, lower panel). Since FtsZ binds to both the N- and the C-terminal domains of MinC, it most likely prevents DNA binding by blocking the binding sites on MinC (and possibly MinD). We confirmed that FtsZ interferes with the DNA binding by the MinCD complex using EMSA (Fig. 3c). Already two minutes of incubation were sufficient to prevent all binding.

### The linker connecting MinC N- and C-terminal domains is needed for DNA binding with MinD and affects the activity of MinC towards FtsZ

To investigate the physiological role of MinCD–DNA binding, we aimed to identify mutants specifically impaired in this activity while still capable of supporting normal mid-cell Z-ring placement. This approach would allow us to rule out indirect effects caused by general cell division or morphological defects.

Considering that MinD has to perform more activities than MinC (interact with the membrane, MinC, and MinE, and hydrolyze ATP to oscillate from pole to pole), we decided to focus on MinC. MinC is, as mentioned above, composed of two domains separated by a linker (Fig. 4a). We decided to test the role of this linker in DNA binding. We predicted that it would not be important for the binding to MinD, since this happens via MinC^C28^. We expected it, however, to play a role in MinC’s activity towards FtsZ, since both MinC domains destabilize FtsZ polymers and their correct positioning towards FtsZ might be important^24, 32^. The linker region of MinC is always referred to as “flexible”, because attempts to crystallize full-length *E. coli* MinC failed so far and, for *T. maritima* MinC, two orientations of MinC^N^ were found in the crystals, which was taken as a proof of the flexibility of the linker^25^. Looking at two available crystal structures of MinC N and C-terminal domains, we found that, in the isolated MinC^N^, only residues 1-101 are resolved^31^, and that, in the isolated MinC^C^, residues 121-231 are resolved^53^. This suggests that the linker region comprises amino acids 102-120 (Fig. 4b). Interestingly, the linker is composed of several amino acids that are found in rigid linkers such as alanine, lysine, and proline^54^; indeed, it is predicted to be rigid by the MEDUSA web server^55^. Nonetheless, given the known flexibility between MinC^N^ and MinC^C^ afforded by the linker, we decided to substitute it with flexible linkers made of various repeats of glycine and serine. We cloned and purified the following MinC linker mutants: GS, 1x AGGSG, 2x AGGSG, 3x AGGSG, 4x AGGSG, and 19RL (Fig. 4b). In the GS construct, MinC^N^ and MinC^C^ are separated by two amino acids, namely glycine (G) and serine (S). In the 1x AGGSG construct (from now on, 1x), the linker region comprises five amino acids, namely alanine (A), glycine (G), glycine (G), serine (S) and glycine (G). In the 2x mutant, this sequence is repeated twice, and so on. The linker in the 19RL mutant is identical to that in the 4x mutant, but lacks the last glycine. Therefore, this mutant linker is as long as MinC’s native linker (that is, it is 19 amino acid long). We then performed EMSAs with wild-type and mutant MinC proteins. We found that all linker variants lost their ability to bind DNA with MinD *in vitro* (Fig. 4c). To investigate the importance of the length of the linker without changing its amino acid sequence, we also constructed mutants of MinC carrying truncations of the linker (Fig. 4d). These mutant MinC proteins were also unable to bind DNA with MinD *in vitro* (Fig. 4e). The linker seems, therefore, crucial for DNA binding. To test the role of the linker further, we investigated the DNA binding ability of the isolated MinC^C^ containing the linker (Fig. 4f, upper panel). This protein bound DNA with MinD even better than full-length MinC (Fig. 4f, lower panel).

**Fig 4.**
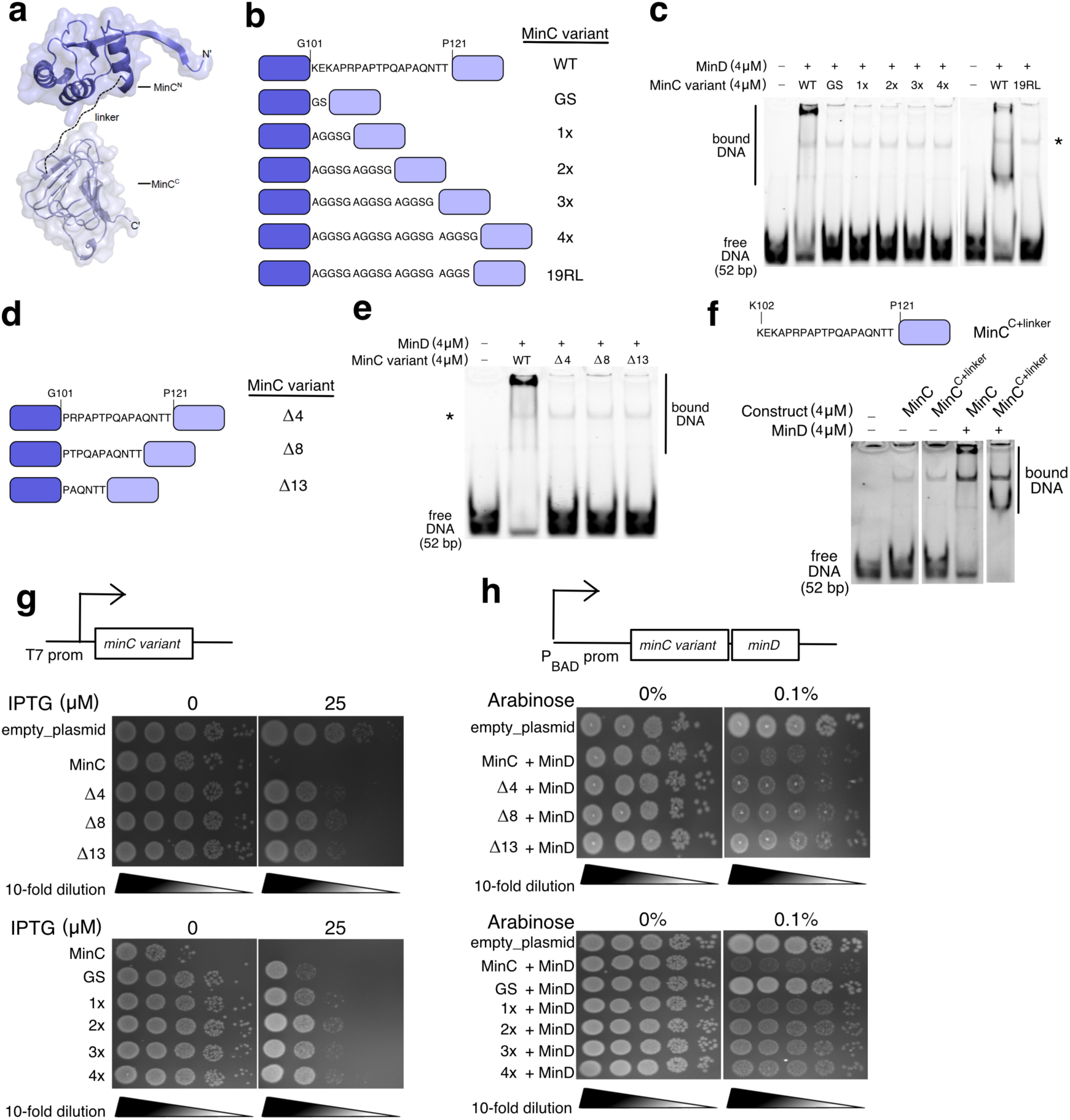
MinC linker is needed for DNA binding with MinD. **a,** Modelled structure of *E. coli* MinC. Crystal structures of MinC N-terminal (PDB ID 4l1c) and C-terminal (PDB ID 5xdm) domains were obtained from the protein data bank (https://www.rcsb.org/). Linker, MinC linker region represented with a dashed line. **b,d**,**f** (upper panel), Schematics of the constructs. **c,e**,**f** (lower panel), Electrophoretic mobility shift assays (EMSAs) performed with the indicated proteins at the indicated concentrations, and HEX-labelled DNA of the indicated length. All reactions were performed in the presence of 1 mM ATP. **c**,**e**, Asterisk indicates the height at which a band is visible, which is observed when incubating the DNA with MinD or MinC alone and does not represent binding by the MinCD complex. **f**, lower panel, White lines indicate unrelated bands that were deleted from the image. The full image is shown in the Supplementary Fig. 5. **g,h**, Upper panel, Schematic of the promoter used in the experiment. Lower panel, representative image of a plate onto which Rosetta BL21 (**g**) or MG1655ΔminB (**h**) cells transformed with the indicated constructs were spotted in a serial dilution. The experiment was repeated three times with similar results.

We confirmed that all the MinC linker mutants were not affected in their ability to bind MinD and be recruited to the liposomes (Fig. S3). For the flexible linker mutants (1x-4x), we additionally checked that they could be dislodged off MinD by MinE (Fig. S4). These results corroborate our assumption that the linker is not involved in the binding of MinC to MinD and MinE.

Finally, we investigated the ability of the linker mutants and truncations to destabilize FtsZ polymers using the spot assay. We expressed either MinC (WT, linker truncations and mutants) alone under the strong IPTG-inducible T7 promoter in *E. coli* Rosetta cells (Fig. 4g, upper panel), or together with MinD under the arabinose-inducible P_BAD_ promoter in MG1655ΔminB cells (Fig. 4h, upper panel). When overexpressed alone, all the linker variants were impaired in their interaction with FtsZ (Fig. 4g, lower panel). In the presence of MinD, the activity towards FstZ improved for all but the GS linker mutant, whose overexpression did not cause filamentation and consequently cell death (Fig. 4h, lower panel). Despite being somewhat affected in their activity towards FtsZ, MinC linker mutants are able to block Z-ring formation when overexpressed with MinD (leading to cell filamentation and cell death), thus they represent a viable option to investigate the impact of DNA binding on chromosomes in living cells for which Z-ring positioning by the Min system is likely in place.

### *E. coli* strains expressing MinC linker mutants from the endogenous locus show chromosomal defects

To construct strains expressing MinC linker mutants at endogenous levels, we applied a genome engineering method based on the CRISPR/Cas9 and the Lambda Red recombination systems^47^, which allowed us to swap the native linker with the 1x-4x variants directly at the endogenous locus. We used two donor strains: MG1655 and MG1655 where a *gfp* gene is inserted in frame downstream of endogenous *hupA* (Fishov strain)^48^. This strain, therefore, expresses the fusion of the alpha subunit of HU (HUα) with GFP and allows visualization of the nucleoid without the use of further stains (e.g. DAPI). Since the procedure itself might affect the cells, we additionally constructed a silent mutant of MinC in both genetic backgrounds. In these silent mutant strains, the nucleotide sequence of the linker was mutated with few alternative codons for the same amino acids. Consequently, they both express wild-type MinC. As a first characterization of the strains, we measured growth curves in LB and M9 media. We found that all mutant strains, including those expressing the silent mutant, grew less than the wild type (Fig. 5a). The 1x linker mutant grew less than all other mutants, for both strains and in both media. Nonetheless, the difference in growth was very moderate.

**Fig 5.**
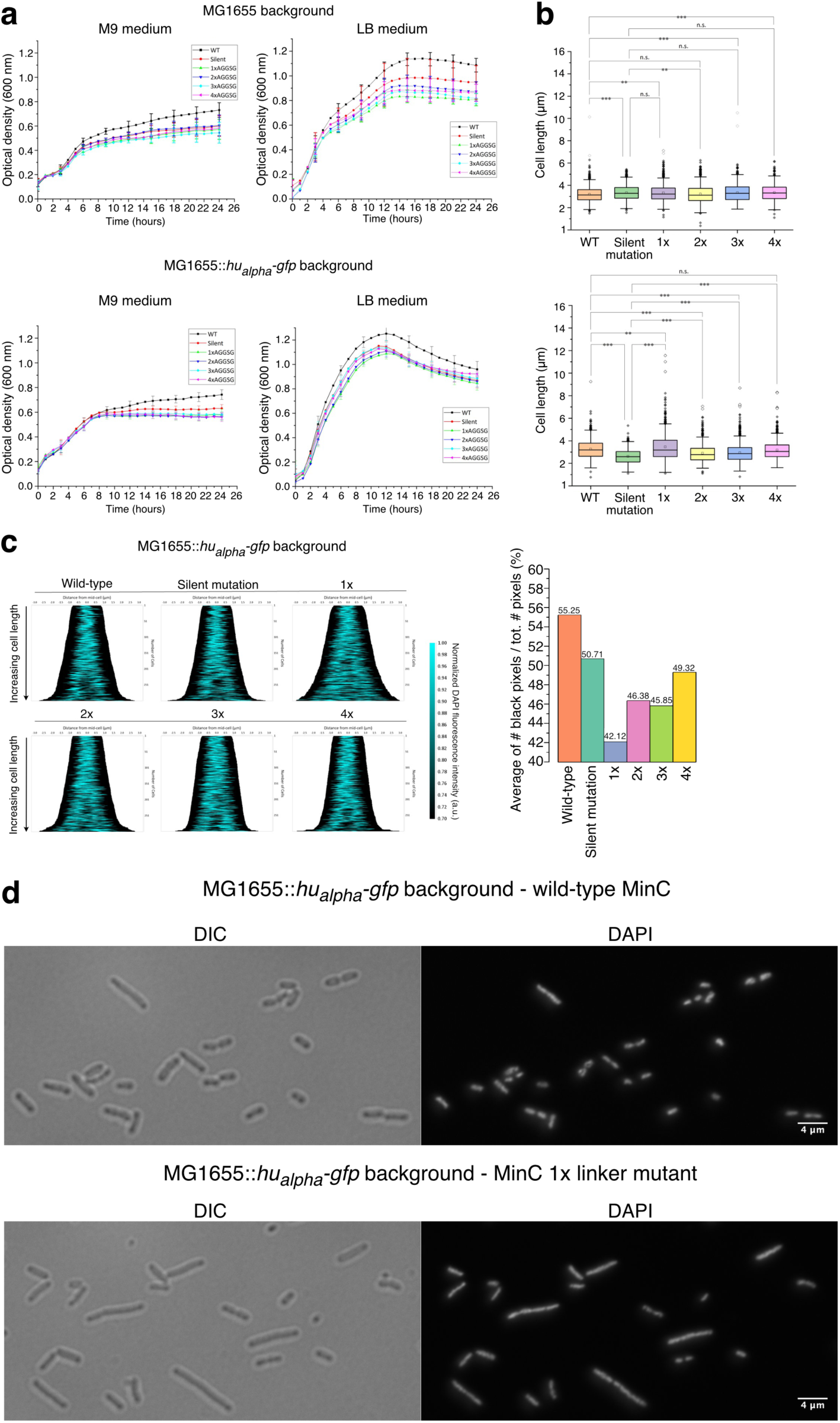
*E. coli* strains expressing MinC linker mutants from the endogenous locus show chromosomal defects. **a,** Growth curves of the indicated strains in the indicated media. **b**, Graphs showing the quantification of the cell length in the indicated strains. More than 500 individual cells were analyzed using the BacStalk software^48^. Shown is the median with the first and third quartiles, as well as outliers. n.s., non-significant. *, p-value < 0.5. **, p-value < 0.01. ***, p-value < 0.001. The p-value was calculated using the one-way ANOVA test. **c,** Left panel, Demographs of the DAPI signal measured via fluorescence microscopy in the indicated strains created with the MicrobeJ plugin of ImageJ. Right panel, Bar graph showing the average of the ratio between the number of black pixels and the total number of pixels in the cell. **a-c**, Silent mutation, strain carrying four silent mutations in the linker region of MinC at the chromosomal locus. **d**, Representative microscopy images of the indicated strains showing the DIC and the DAPI channels.

We then measured cell length using the automated software BacStalk^56^. Note that, in this measurement, minicells were explicitly excluded. We found a statistically significant difference between the wild type and almost each mutant, with the exception of the 2x linker mutant in the MG1655 background and the 4x linker mutant in the Fishov background (Fig. 5b). In the MG1655 background, the median was almost identical in all strains (Fig. 5b, upper panel). In the Fishov background, the silent mutant, as well as the 2x and 3x mutants gave rise to smaller cells than the wild type (Fig. 5b, lower panel). The 1x mutant, despite having a similar median as the wild type, had many longer cells.

We then employed fluorescence microscopy using DAPI to stain the nucleoid and constructed demographs of the DAPI signal (Fig. 5c and Fig. S6). Note that, albeit the Fishov strains express HUα-GFP, which allows visualization of the chromosome in live cells, we opted for fixing and staining the cells with DAPI to treat all samples in the same way. Observation of the demographs with the naked eye revealed clear differences for several of the mutants in the Fishov background (Fig. 5c, left panel). To resolve defects not obvious to the naked eye, we decided to calculate a measure that would indicate the spread of the DAPI signal in the cell. For less well-segregated chromosomes or less compacted chromosomes, we expect a higher spread. We therefore calculated the ratio between the number of black pixels (which indicate the area of the cell without DNA) and the total number of pixels in the cell. We calculated the average of this ratio for all cells and expressed it as a percentage (Fig. 5c and Fig. S6, right panels). With this measure, it is possible to appreciate that, in the Fishov background, chromosomes are more spread for all the MinC mutants, interestingly even for the silent one (Fig. 5c, right panel). The clear phenotype of the 1x linker mutant can be appreciated from fluorescence microscopy images (Fig. 5d).

## Discussion

The role of the Min system in mid-cell determination and Z-ring positioning is well established. Years ago, the Min system was proposed to also contribute to chromosome segregation^41^, but many details have remained unclear. Here we show that the MinCD complex binds DNA more strongly than either protein alone. MinE dissociates the MinCD-DNA complex, which would be important to ensures that the DNA binding is transient in the cellular context. Moreover, we used CRISPR/Cas9 coupled with the Lambda Red recombination to construct strains that are identical to the parental ones except that endogeous MinC carries a modified linker, which we demonstrated to abrogate DNA binding *in vitro*. Some of these mutant strains have chromosomes that are more spread throughtout the cell when the alpha subunit of the histone-like protein HU is expressed endogenously tagged to GFP. This could be due to the fact that HU plays a role in maitaining the correct structure of the chromosome and the fusion to GFP may not be as functional as the wild-type and therefore this strain may already suffer from mild defects in chromosome structure/compaction or other processes related to HU^57–60^. Indeed, comparison of demographs for wild-type strains shows better separation in longer cells in MG1655 than in MG1655::*huα-gfp* (Fig. 5c and Fig. S6, left panels). These mild defects are exacerbated in the MinC linker mutant strains.

As mentioned already, the mutant strains in the MG1655 background show only very minor defects. While the DNA binding activity of the mutants was assessed *in vitro*, translating these results to the cellular context is challenging due to the complexity of cellular environments, including varying protein concentrations and additional factors. It is possible that the mutations only partially impair DNA binding in the cell, leading to milder defects than observed *in vitro*. From this analysis, it is not possible to discern if the effects we observed are due to chromosome segregation defects, less DNA compaction, or a combination of both.

It is difficult to entirely disentagle the activity of the Min system towards FstZ and in chromosome segregation/organization because, if altered, both processes lead to the formation of minicells. There are several reports stating the presence of minicells as a consequence of failed chromosome segregation in various bacterial species^61–65^. In *E. coli*, *muK* mutants have been reported to give rise to anucleate, but normally sized cells^66^; however, microscopy images clearly show the presence of minicells (see Fig. 2 in ref^66^). By taking snapshots of all the constructed strains in the microscope and quantifying the percentage of minicells formed, we found that, in the MG1655 background, all the linker mutants produced minicells compared to the wild-type strain and the strain expressing the silent mutant, with the 2x linker mutant producing the highest amount of minicells, followed closely by the 3x mutant (Fig. S7). In the Fishov backround, the percentage of minicells produced by the linker mutants is overall higher (Fig. S8).

We were surprised to find differences between wild-type strains and those with silent mutations in MinC linker. We excluded the possibility that these differences are due to protein levels (Fig. S9). To understand if they might be due to off-target mutations introduced by Cas9, we sequenced the whole genome of all Fishov strains, including the wild-type and the silent mutant. We surprisingly found the same two mutations – in the *sspA* and *mepA* genes – in all strains that underwent the CRISPR/Cas9+Lambda red recombineering procedure. The 4x linker mutant has one additional mutation in the *EGEAPM_04336* gene. Using PCR, we checked if the same mutations are present in the MG1655 mutant strains. This is not the case. To understand the reason behind this strange phenomen, we would need to repeat the genome engineering procedure to see if the same mutations arise again and only in the Fishov backgroud. It cannot be excluded that the mutations in SspA and MepA are responsible for the phenotype seen in the silent mutant strain. Given the difference between the silent and the other mutants, though, we can conclude that the MinC linker mutants have additional effects on the chromosomes.

The electron microscopy results reveal that DNA can associate with MinCD co-polymers (Fig. 2). With free dsDNA fragments, we observed sheet-like structures whose widths scaled with DNA length, consistent with DNA running perpendicular or at an angle to the filaments. In contrast, dsDNA fiducials attached to gold beads aligned along single filaments without bridging neighboring ones. The linear arrangement of the beads closely followed the curved filaments, rarely switching between them, and neighboring filaments were only sparsely labeled. At present, we cannot explain the different behavior of free DNA fragments versus fiducials, nor the strikingly regular bead spacing. One possible contributing factor is electrostatic repulsion between DNA-coated beads.

The fiducial data suggest cooperativity: binding of an initial bead may promote the recruitment of additional beads to the same filament, producing bead “lines.” Individual bead-binding events appear rare. Such cooperative binding, if it occurs *in vivo*, could help MinCD filaments interact more stably with extended DNA regions, potentially stabilizing replicated chromosomes near the poles. Although the existence of MinCD co-polymers in living cells is debated^67^, it is possible that they exist.

The data presented in this work are not sufficient to conclude a role for MinCD in chromosome segregation, and further experiments are required. Future work might address whether MinCD can also bind ssDNA or RNA, and whether DNA binding could modulate Min oscillation dynamics, for example by slowing oscillations differently depending on DNA content and chromosome positioning.

## Materials and methods

### Strains and growth conditions

The strains used in this study are listed in Supplementary Table 1. The *E. coli* TOP10 strain was used for cloning, and the *E. coli* Rosetta™(DE3)pLysS (Novagen) strain was used for overproduction and protein purification. All *E. coli* strains were grown in LB or M9 + 0.4% glucose supplemented with the appropriate antibiotic(s) at 37 °C with shaking.

### Construction of plasmids

pET28a-MinD and pET28a-MinE plasmids were constructed as previously described^41, 68^. The *minC* gene was amplified from *E. coli* MG1655 genomic DNA using *BamHI*_MinC_FP and *NotI*_MinC_RP primers. The amplicon and pET28a empty plasmid were double digested using *BamHI* and *NotI* and then ligated to get pET28a-MinC. Similarly, the *ftsZ* gene was amplified from *E. coli* MG1655 genomic DNA using *BamHI*_FtsZ_FP and *HindIII*_FtsZ_RP primers. The amplicon and pET28a empty plasmid were double digested using *BamHI* and *HindIII* and then ligated to get pET28a-FtsZ. pET28a-MinC-GS was obtained by amplifying the whole pET28a-MinC plasmid except the linker region (i.e., residues 102 to 120) using phosphorylated primers MinC_GS_FP and MinC101_RP with the forward primer containing the codons for “GS”, and then self-circularizing the linear PCR product using T4 DNA ligase. Using the same strategy and same reverse primer (i.e., MinC101_RP), pET28a-MinC-1xAGGSG (using primer MinC_1xAGGSG_FP), pET28a-MinC-2xAGGSG (using primer MinC_2xAGGSG_FP), pET28a-MinC-3xAGGSG (using primer MinC_3xAGGSG_FP), pET28a-MinC-4xAGGSG (using primer MinC_4xAGGSG_FP), pET28a-MinC19RL (using primer MinC_19RL_FP), pET28a-MinC-Δ4 (using primer MinC_del4_FP), pET28a-MinC-Δ8 (using primer MinC_del8_FP) and pET28a-MinC-Δ13 (using primer MinC_del13_FP) were constructed. Also, using the same strategy, pET28a-MinC^C+linker^ (using primers MinC102_FP and pET28a_*BamHI*_RP) was constructed.

pBAD33-MinCD was obtained from a previously constructed pBAD33-MinCDE^68^ by removing the *minE* gene by linearization and self-circularization using phosphorylated primers pBAD33_*HindIII*_FP and MinD_RP. Using the same strategy and primers mentioned above for pET28a-MinC linker variants, pBAD33-MinCD-GS, pBAD33-MinCD-1xAGGSG, pBAD33-MinCD-2xAGGSG, pBAD33-MinCD-3xAGGSG, pBAD33-MinCD-4xAGGSG, pBAD33-MinCD-Δ4, pBAD33-MinCD-Δ8 and pBAD33-MinCD-Δ13 were constructed.

For all the PCR amplifications, high-fidelity Phusion polymerase (New England Biolabs) was used. Plasmids were amplified by transforming them into chemically competent *E. coli* TOP10 cells (Invitrogen™) and were isolated using the QIAprep® Spin Miniprep Kit (Qiagen) as per the manufacturer’s protocol. The constructs were validated via Sanger sequencing (GATC Biotech, Germany). The list of primers used in this study is given in Supplementary Table 2.

### Constructions of strains with MinC linker variants

To generate strains expressing MinC linker mutants from the endogenous locus we employed a method combining CRISPR/Cas9 and Lambda red homologous recombination^47^. The two plasmids, pcas9 and pTarget (Lambda Red Homologous Recombination Plasmid and CRISPR HR gRNA Plasmid), were purchased from Sigma-Aldrich. pcas9 was transformed into wild-type *E. coli* MG1655 and the cells were plated on LB agar plates with 50 μg/mL kanamycin. The strain harbouring pcas9 was cultured overnight in LB medium at 30°C with shaking (250 r.p.m.). The next day, a fresh culture was started from the overnight culture using 3 mL fresh LB medium with 50 μg/mL kanamycin. The culture was then grown at 30°C until the OD_600_ reached 0.3-0.5. 1.5% arabinose was used to induce the expression of the Lambda red recombination proteins for 1 hour. Cells were then collected and washed twice with 40 μL ice-cold 20% glycerol to make them become electrocompetent. Donor DNA and pTarget were transformed into the cells by electroporation. Cells bearing both pcas9 and pTarget were selected on LB agar plates with 50 μg/mL kanamycin and 100μg/mL ampicillin. The donor DNA was obtained by PCR using high-fidelity Phusion polymerase (ThermoFisher). After electroporation, the cells were plated on LB agar plates with 50 μg/mL kanamycin and 100μg/mL ampicillin. A randomly selected colony for each strain was picked from the plate and a PCR was performed to check for the presence of the mutations. This colony was grown overnight and then a diluted culture was spread on a LB agar plate with 5% saccharose. The plate was left overnight at 37 °C. The following morning, eight colonies for each strain were randomly selected and streaked onto three plates: a LB agar plate with no antibiotic, a LB agar plate with 100 μg/mL ampicillin, and a LB agar plate with 50 μg/mL kanamycin. The plates were incubated overnight at 37 °C. The next day, the colonies that died on the plates with the antibiotics were considered cured of pCas9 and pTarget and were therefore selected by using the LB plate without antibiotic. The correctness of the sequence of every MinC linker mutant was checked by colony PCR followed by Sanger sequencing.

### Protein purification

Protein purifications were performed as previously described^41^ with minor modifications. Rosetta cells carrying the desired plasmid were grown to an OD600 of 0.5-0.9 and induced with 1 mM IPTG for 3 hours at 37 °C. Cells were collected by centrifugation (4.000-5.000 r.p.m.) at 4 °C for 20 minutes and stored at −20 °C. Pellets were thawed on ice and resuspended in lysis buffer. The buffers used in this study are listed in Supplementary Table 3. Cells were then lysed by sonication and the crude lysate was clarified by centrifugation (20.000 r.p.m.) for 20 min at 4 °C. Proteins were purified using the Profinia™ Protein Purification System (Bio-Rad), with a BIO-RAD® Bio-Scale™ Mini Profinity™ IMAC Cartridge (1 mL) and the BIO-RAD® Bio-Scale™ Mini Bio-Gel® P-6 Desalting Cartridge (10 mL). The column was washed with imidazole buffer and the protein was eluted with elution buffer, followed by exchange of the elution buffer for storage buffer. The purified proteins were stored in small aliquots at −80 °C. Protein concentrations were estimated based on the absorbance at 280 nm with a NanoDrop spectrophotometer (Thermo Scientific). Proteins were cleared from any aggregates by centrifugation (21.000 g) for 30 min at 4 °C prior to use.

### Elimination of the His tag

His tag cleavage was performed for the other aliquot by incubating the proteins with thrombin agarose beads (Thrombin CleanCleave™ Kit from Sigma-Aldrich) for 3 hours at room temperature on a rotating wheel. After the His tag removal, the proteins were stored at −80 °C in 50 μL aliquots.

### Electrophoretic mobility shift assays

Proteins were incubated with 50 nM 5’ HEX-labelled (hexachloro-6-carboxy-fluoresceine) DNA in a buffer containing 50 mM HEPES-KOH pH 7.25, 150 mM KCl, 10 % glycerol, 0.1 mM EDTA pH 8.0, 1 mM ADP/ATP and 5 mM MgCl_2_ at room temperature for 10 min. The reaction mixture was then loaded on a 6 % native-PAGE gel. The gel was run in 0.5x TBE buffer for 20 min at constant voltage (150 V). Finally, the gel image was obtained using an Amersham Typhoon Gel and Blot Imaging Systems (GE Healthcare).

### Preparation of double-stranded DNA probes for EMSAs

Two complementary oligos were mixed 1:1 in NEB buffer 2 and the mixture was heated up to 98°C in a heat block. Only one of the oligos was HEX-labeled at the 5’ end. After 2 minutes, the heat block was switched off, and the samples were left to gradually cool down. DNA probes longer than 120 bp were amplified via PCR using a forward primer with a HEX label, followed by gel purification of the fragment using the Qiagen PCR clean up kit.

### Liposome preparation

*E. coli* total lipids (powder) were purchased from Avanti Polar Lipids. The lipids were dissolved in liposome buffer (20 mM HEPES pH 7.5, 150 mM NaCl and 5 mM β-mercaptoethanol) to a final concentration of 20 mg/mL and left for 30 minutes at 37 °C. For better hydration, the sample was frozen in liquid nitrogen and heated up to 37 °C in a heat block a total of 5 times. Lipid vesicles were obtained using the Avanti Mini Extruser with a 200 nm filter. The sample was passed through the filter 20 times, aliquoted and stored at −80 °C.

### Liposome co-sedimentation assay

We used a previously described protocol^41^. Liposomes (320 μg/mL in Figs. 1d and 3 c,d, 500 μg/mL in Supplementary Fig. 3 and 160 μg/mL in Supplementary Fig. 4) were incubated for 10 min with the proteins (6 μM MinC and MinD in Figs. 1d and 3 c,d, 2 μM MinC and MinD in Supplementary Fig. 3, and 1 μM MinC and MinD in Supplementary Fig. 4) and 1 mM of either ADP or ATP in 50 μL reaction volume in ATPase buffer (25 mM Tris-HCl, pH 7.5, 50 mM KCl, 5 mM MgCl2, 5% glycerol). The reactions were pelleted by centrifugation at 21.000 g for 15-30 min, the pellets were resuspended in 50 μL ATPase buffer, and supernatant and pellet samples were analyzed by SDS-PAGE, followed by Coomassie staining to detect the proteins.

### Liposome co-sedimentation in the presence of plasmid DNA

Purified proteins (varying concentrations) were incubated with 800 μg/mL liposomes, 32 ng/ μL plasmid DNA (4.7 kb) and 1 mM ADP/ATP in storage buffer supplemented with 5 mM MgCl_2_. After 15 minutes of incubation at room temperature, the reaction mixture was spun down in a table top centrifuge at 21.000 g for 20 minutes. The supernatant was removed and mixed in a 1:1 ratio with 2x Laemli buffer. The pellet was dissolved in 1x Laemli buffer with a volume matching that of the supernatant. After heating the samples for 5 minutes at 95 °C, the protein content of pellet and supernatant fractions was analysed with SDS-page followed by Coomassie staining. A 1% agarose gel stained with Gel Red™ Nucleic Acid Stain was used to analyse the DNA content. The intensity of the DNA and protein bands was quantified with ImageJ.

### Negative-stain transmission electron microscopy

The filaments were formed by incubating MinC (4 µM) with MinD (4 µM), 1 mM ATP, 10 mM MgCl2 and DNA (50 nM) for 15 minutes on ice. For the experiment shown in Fig. S2, the DNA was omitted. Grids with continuous carbon were glow discharged and 3 µl of sample were pipetted on the grid. After 30 seconds, the liquid was removed and 10 µl of 1% uranyl acetate in water were pipetted on the grid and incubated for 30 seconds. The excess stain was removed and the grids were dried. For each condition 25 images were randomly taken and the number of filaments in the sheets determined. Single filaments were not counted.

### Preparation of DNA fiducials

Single-stranded fiducials (ss fiducials) were prepared by coating citrate gold beads of 5 nm (PROVIDER) with thiolated single-stranded dT DNA oligomers. To obtain double-stranded fiducials (ds fiducials), an excess of dA oligomers was added and annealed to the ss fiducials.

### Binding of DNA fiducials to MinCD co-polymers

MinCD co-polymeric filaments were produced by mixing 4 μM of MinD with 4 μM MinC. Larger aggregates were removed by spinning the protein solutions at 15 000 x g for 30 minutes. Polymerization was started by adding ATP (200 µM final concentration) and MgCl_2_ (10 mM final concentration) and incubating the mixture for 10 minutes on ice. An equal volume of the fiducials was added and the 3 μl of the sample was plunge-frozen on Quantifoli grids (0.6/1, 2/1, 3.5/1) with blotting times between 5 and 15 seconds at 100% humidity. Grids were observed in a Krios instrument (Thermo Fisher) and screened at low magnification (135x) to identify good quality regions. Best grids were selected for targeted imaging (Latitude/Gatan) or random imaging (EPU/FEI). For the analysis, 2x binned images were used at a pixel size of 2.74 Å. After manual exclusion of low-quality images, the gold beads were localized (imodfindbeads) and all linear distances computed (imod-dist) with an ad-hoc script (Supplementary Information). Occurring instances smaller than 250 pixels were counted (in 125 bins for smoothing the data) and plotted.

### Heat treatment of ds fiducials

ds fiducials were heated to 95 °C and cooled to 4 °C over 30 minutes.

### Spot assay

Spot assay was performed as previously described^21^. Briefly, the plasmids were transformed into either *E. coli* ROSETTA or MG1655Δ*minB* strain. A colony was picked and suspended in a nutrient broth without any antibiotic. The cells were then serially diluted and spotted on agar plates with different IPTG or arabinose concentration. The plates were then incubated overnight at 37 °C and imaged using a gel imaging system (Analytik Jena).

### Growth curve measurement

Each strain was grown overnight at 37°C and shaken at 200 rpm in the selected medium (either LB or M9 + 0.4% glucose). The cultures were diluted to OD_600_=0.1 and added in triplicates to a 96-well microplate. The OD_600_ of each well was measured every 10 minutes for 24 hours by the CLARIOstar® Plus Microplate Reader from BMG labtech. The microplate was incubated at 37°C and shaken before every measurement. The experiment was repeated three times.

### Cell length measurement

The *E. coli* MG1655 and Fishov strains were grown overnight at 37°C and shaken at 200 rpm in M9 + 0.4% glucose medium. The morning after, cultures were diluted to OD_600_=0.1 and incubated in the same conditions for 4 hours. After reaching an OD_600_ of approximately 0.4, bacteria were fixed with a 3,7 % formaldehyde solution in PBS. 100 nM DAPI were added to each sample prior to microscopy analysis. Microscopy pictures were taken on a Zeiss Axio Observer Z1/7 (Carl Zeiss Microscopy). 3 μL of the bacteria-containing solution were spread on a 1% agarose pad on metal microscope slides. The BacStalk software^56^ was used to detect cells and automatically measure their length. The size and position of nucleiods in the cells were determined with the MicrobeJ plugin of ImageJ by measuring the intensity of the DAPI fluorescence throughout the whole length of the bacteria.

### Demographs

The demographs were created using the MicrobeJ plugin of ImageJ. Cells with a length between 1.5 and 6 µm were selected using the selection parameters of the software. The cells were sorted by cell length in ascending order from top to bottom. Only the pixels with a fluorescence intensity greater than 0.7 were plotted to increase the readability of the graphs.

### Sample preparation for mass spectrometry

An overnight culture of each mutant strain, plus the parental strains, was grown in LB at 37°C and shaken at 200 r.p.m. The morning after, a fresh culture of 2 mL was started with OD_600_=0.1, and let grown at 37 °C and shaken at 200 r.p.m. for two hours. The cultures were then pelleted at maximal speed in a table-top centrifuge. The pellet was resuspended in 4x Laemli buffer diluted in PBS (to obtain 1x Laemli buffer) and boiled at 98 °C for ten minutes. The samples were then loaded on a SDS gel, which, after the run, was stained with Coomassie and sent over for mass spectrometry analysis.

### Mass spectrometry

#### In-gel tryptic digestion

Upon SDS-PAGE, Coomassie-stained bands were manually excised from the gel and processed as described before^69^. Two gel slices covering theoretical molecular weight from ± 18 kDa up to 37 kDa were collected per gel lane. The extracted peptides from each gel slice were analyzed using LC-MS/MS, generating separate raw data files for each slice.

### LC-MS/MS analysis

Nanoflow LC-MS^2^ analysis was performed with an Ultimate 3000 liquid chromatography system coupled to an Orbitrap QE HF (Thermo Fisher). An in-house packed analytical column (75 µm x 200 mm, 1.9 µm ReprosilPur-AQ 120 C18 material (Dr. Maisch, Germany) was used. Mobile phase solutions were prepared as follows, solvent A: 0.1% formic acid / 1% acetonitrile, solvent B: 0.1% formic acid, 89.9% acetonitrile. Peptides were separated in a 60 min linear gradient started from 3% B and increased to 23% B over 50 min and to 38% B over 10 min, followed by washout with 95% B. The mass spectrometer was operated in data-dependent acquisition mode, automatically switching between MS and MS2. MS spectra (m/z 400–1600) were acquired in the Orbitrap at 60,000 (m/z 400) resolution and MS2 spectra were generated for up to 15 precursors with a normalized collision energy of 27 and an isolation width of 1.4 m/z.

### Database search and data processing

The MS/MS spectra were searched against the Swiss-Prot *Saccharomyces cerevisiae* (UP000002311, November 2019) and *Escherichia coli (*UP000000625, November 2019) protein databases and a customized contaminant database (part of MaxQuant, MPI Martinsried) using Proteome Discoverer 2.5 with Sequest HT (Thermo Fisher Scientific). A fragment ion mass tolerance was set to 0.02 Da and a parent ion mass tolerance to 5 ppm. Trypsin was specified as an enzyme. Carbamidomethyl was set as fixed modification of cysteine and oxidation (methionine), and deamidation (asparagine, glutamine) as variable modifications of peptides. Acetylation, methionine-loss, and a combination of acetylation and methionine-loss were set as variable modifications of the protein terminus. For feature linking and mapping, RT tolerance [min] and mass tolerance [ppm] were set to 0.001 and 0.0005, respectively. Peptide quantification was done using a precursor ion quantifier node with the Top N Average method (N=3) set for protein abundance calculation. For each protein, the intensities recorded in the two gel slices were summed to obtain the total protein abundance. This summation approach was applied to account for any distribution of proteins between the gel slices. To account for the measurement inconsistencies, median normalization was applied.

### Statistics

Statistical analyses were performed in Origin Pro.

## Acknowledgments

We thank Victor Sourjik (MPI, Marburg) for critical reading of this manuscript, Luis Serrano (CRG, Barcelona) and Victor Sourjik for discussions during the course of this work, and Chris Van der Does (University of Freiburg) for the generous donation of liposomes. We acknowledge the technical support of Core Facility for Mass Spectrometry and Proteomics of Center for Molecular Biology (ZMBH) of Heidelberg University. Core Facility for Mass Spectrometry and Proteomics is funded by the ZMBH and partially funded by the CellNetworks Core Technology Platform (CCTP) of Heidelberg University. The CCTP is funded in part by the Federal Ministry of Education and Research (BMBF) and the Ministry of Science Baden Württemberg within the framework of the Excellence Strategy of the Federal and State Governments of Germany. Part of this work was performed at the Center for Quantitative Biology (BioQuant), Heidelberg. L.K. and N.P. were funded by the German Research Foundation (DFG) (grant no. VE776/2-1 to B.D.V.).

## Contributions

B.D.V. conceived the study. N.P. and L.K. purified the proteins, performed EMSAs and liposome co-sedimentations. N.P. performed the spot assays. J.L. constructed the mutant strains and prepared the samples for mass spectrometry. M.L. characterized the strains in terms of growth, cell length and chromosome segregation. G.H. and J.Z. performed all the negative-stain transmission microscopy. M.U. provided the DNA fiducials. M.A.Ö. helped with structural analyses. S.M., M.L and T.R. performed mass spectrometry and data analysis. B.D.V. supervised the work of N.P., L.K., M.L. and J.L., provided the funding and wrote the manuscript. All authors commented on and approved of the manuscript.

## Supplementary Figures and Legends

**Fig S1.**
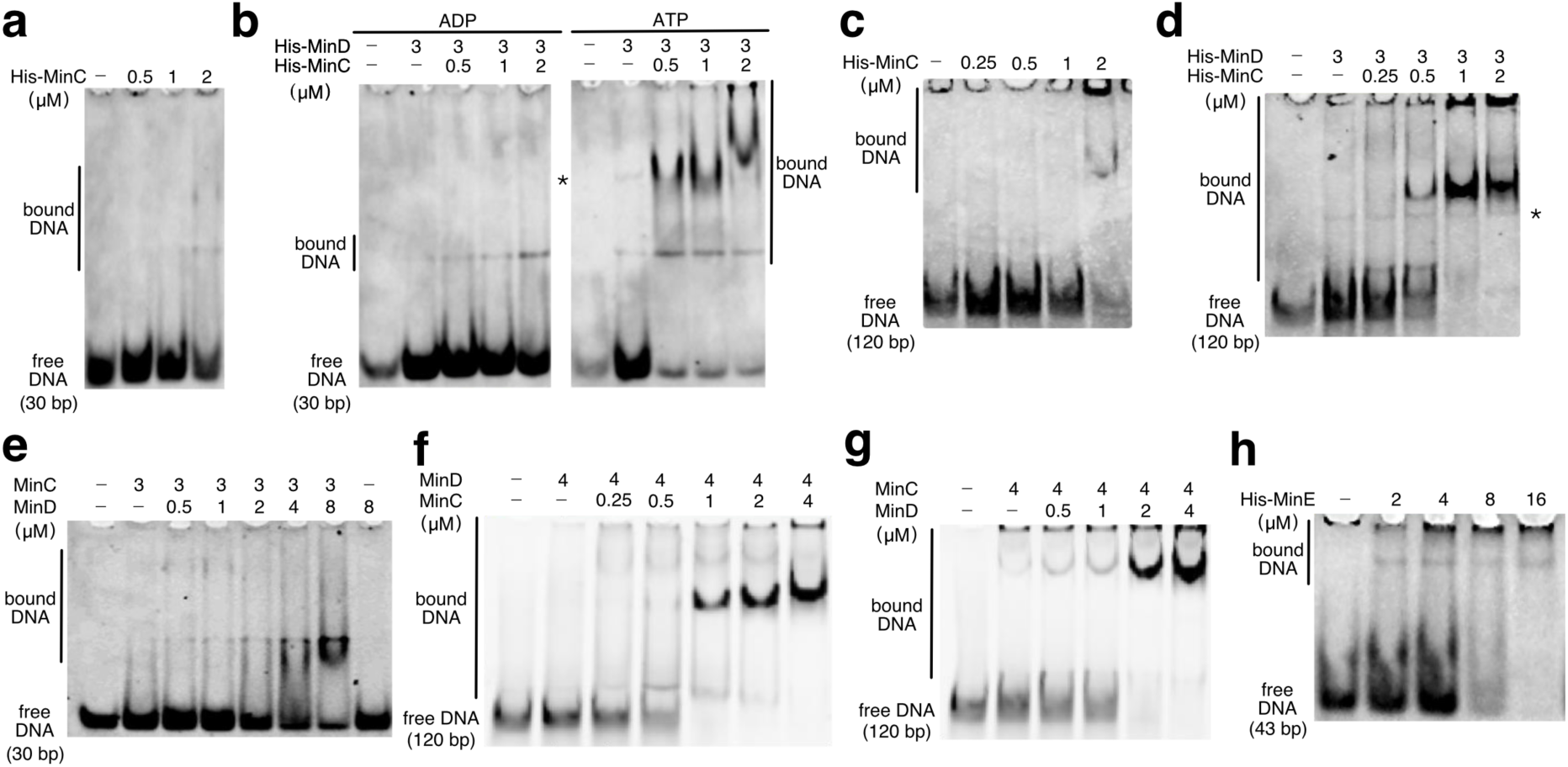
The His-tag enhances the binding of the Min proteins to the DNA. **a-h**, Electrophoretic mobility shift assays (EMSAs) performed with the indicated proteins at the indicated concentrations, and HEX-labelled DNA (50 nM) of the indicated length. **a**, **c-h**, All reactions were performed in the presence of 1 mM ATP. **d,** Asterisk indicates the height at which a band is visible, which is observed when incubating the DNA with MinD alone and does not represent binding by the MinCD complex.

**Fig S2.**
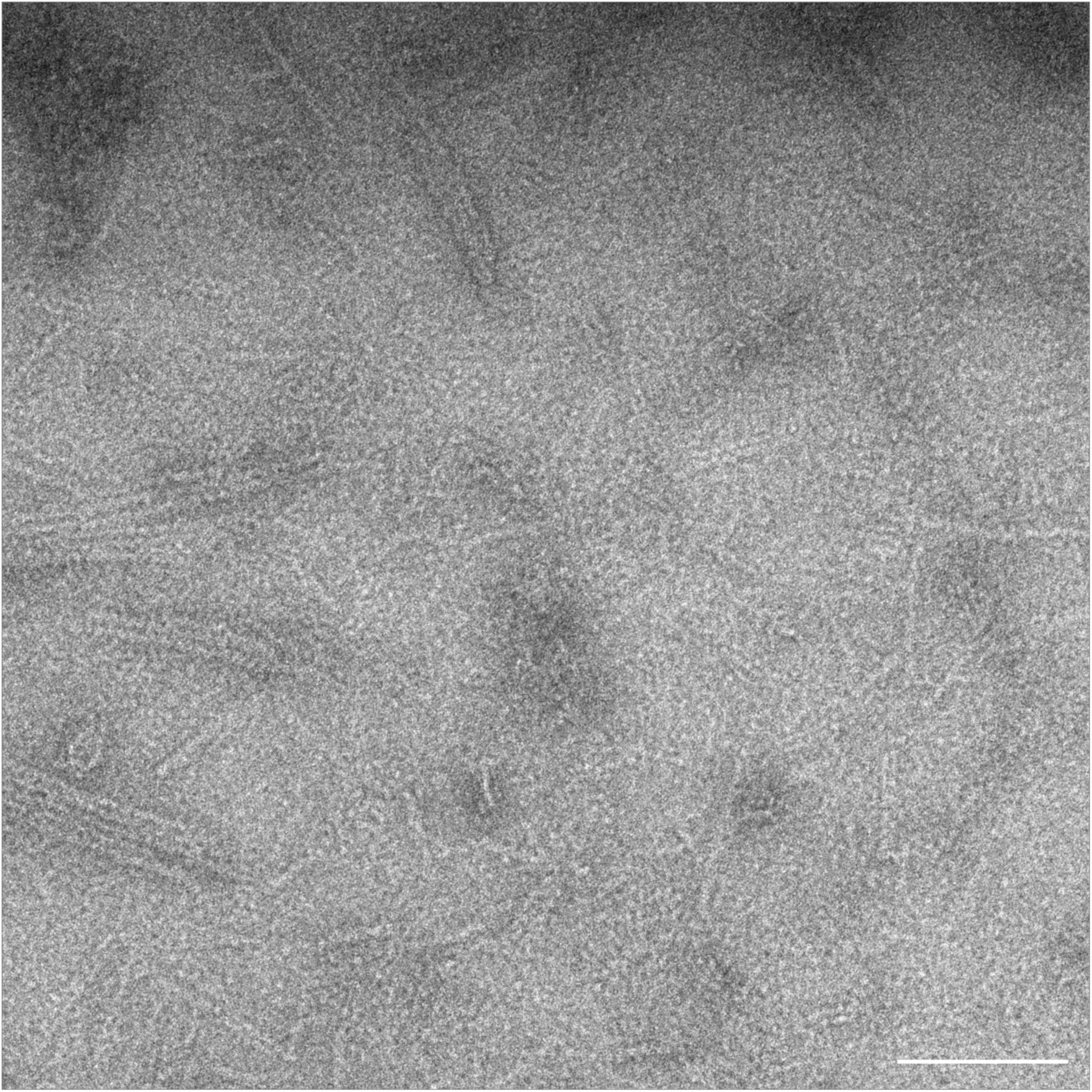
MinC and MinD incubated with ATP occasionally form sheets. Negative-stain electron microscopy image showing the MinCD co-polymers in the absence of added DNA. Scale bar, 100 nm.

**Fig S3.**
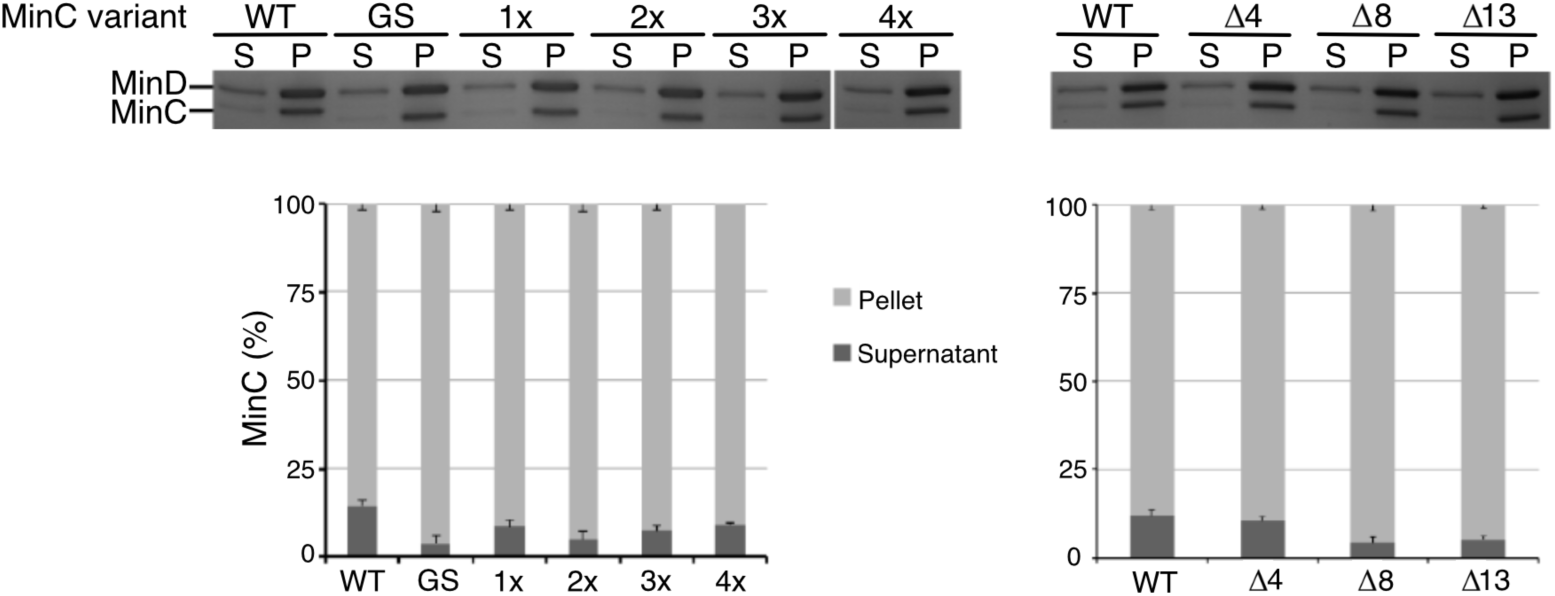
Min linker mutants are recruited to liposomes by MinD. Liposome co-sedimentation assay with MinD and MinC linker variants. MinD (2 μM) was incubated with MinC (2 μM) and 0.5 mg/mL liposomes in a buffer containing 1 mM ATP. Upper panel, representative Coomassie-stained SDS-PAGE gels showing MinC and MinD in the supernatant (S) and pellet (P) fraction. White line indicates that the 4x sample was run in another gel compared with the other samples. Lower panel, Bar graph showing the quantification of MinC in the supernatant and pellet. Data represent mean ± S.E.M of n=3 independent experiments.

**Fig. S4.**
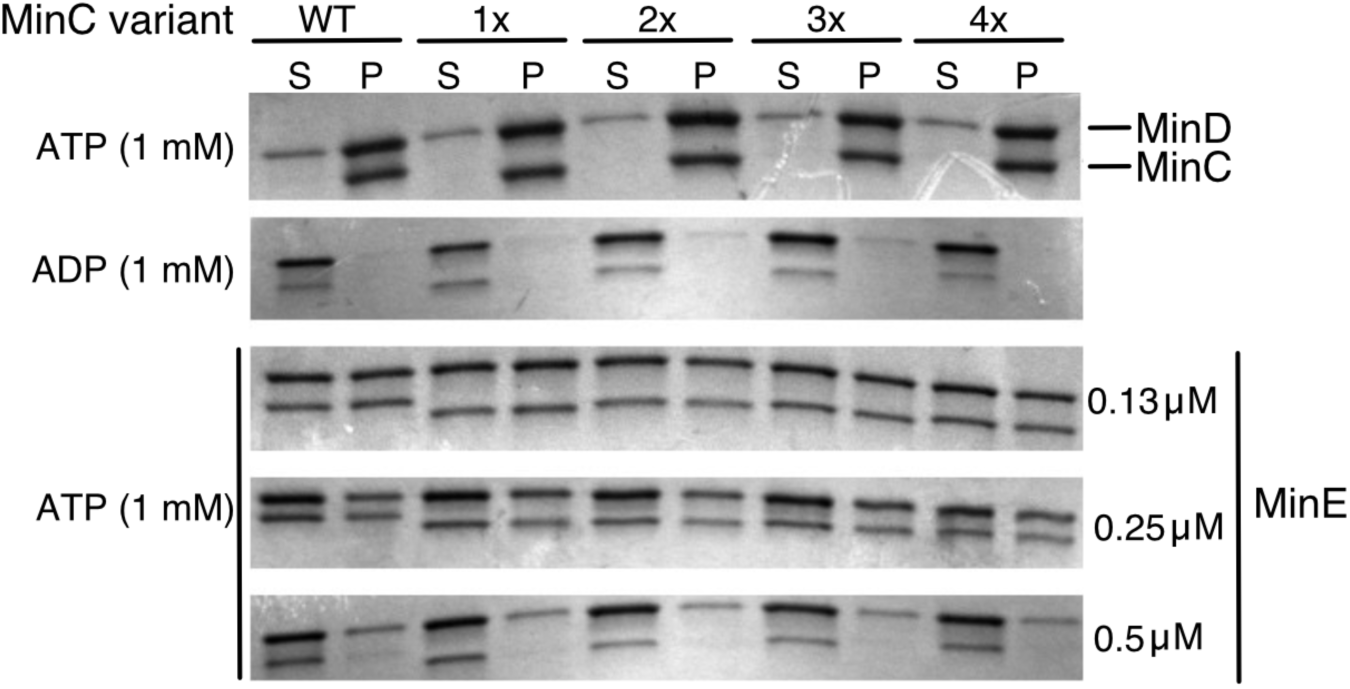
MinC linker mutants are dislodged off MinD by MinE. Liposome co-sedimentation assay with MinD, MinC linker variants and MinE. MinD (1 μM) was incubated with MinC (1 μM), the indicated amounts of MinE and 160 μg/mL liposomes in a buffer containing 1 mM ATP. Representative Coomassie-stained SDS-PAGE gels showing MinC and MinD in the supernatant (S) and pellet (P) fraction. The experiment was repeated three times with similar results.

**Fig. S5.**
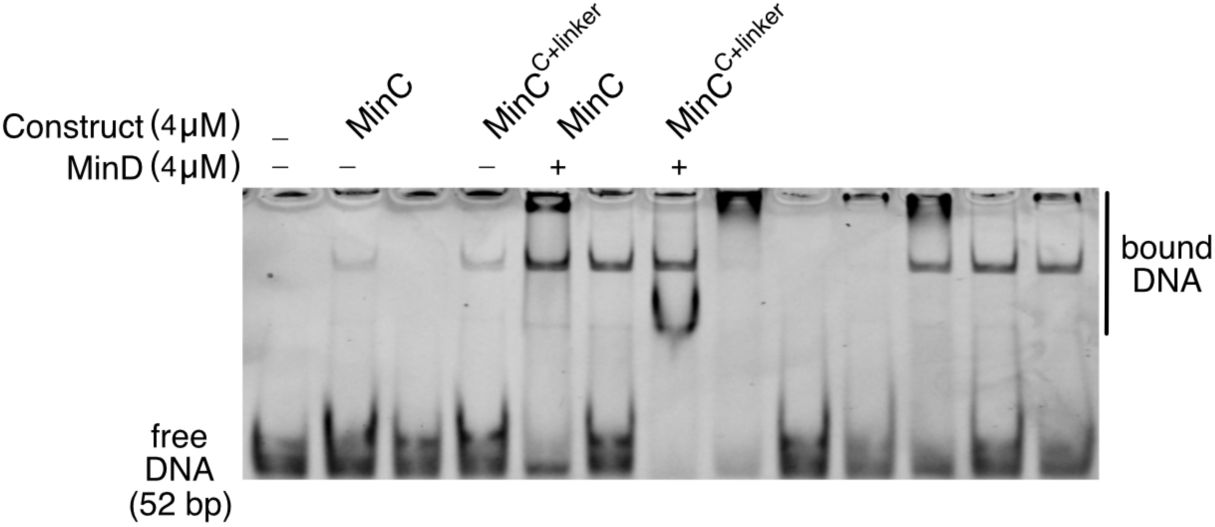
Full image of the EMSA shown in Fig. 4c. Only the relevant bands are labeled. In the other bands, other constructs which are not discussed in this study were tested.

**Fig. S6.**
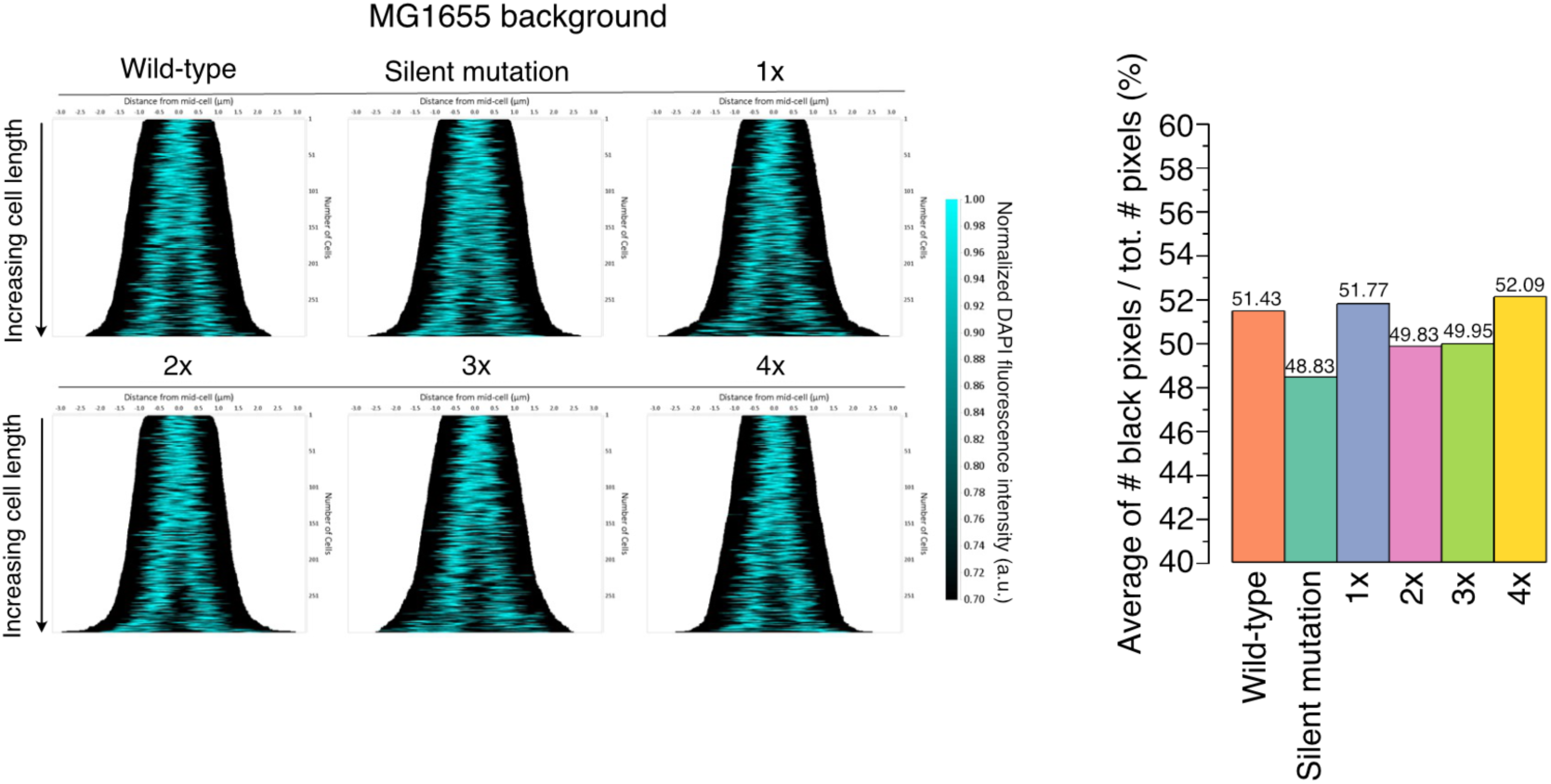
MinC linker mutants show only minor chromosomal defects in the MG1655 background. Left panel, Demographs of the DAPI signal measured via fluorescence microscopy in the indicated strains created with the MicrobeJ plugin of ImageJ. Right panel, Bar graph showing the average of the ratio between the number of black pixels and the total number of pixels in the cell.

**Fig. S7.**
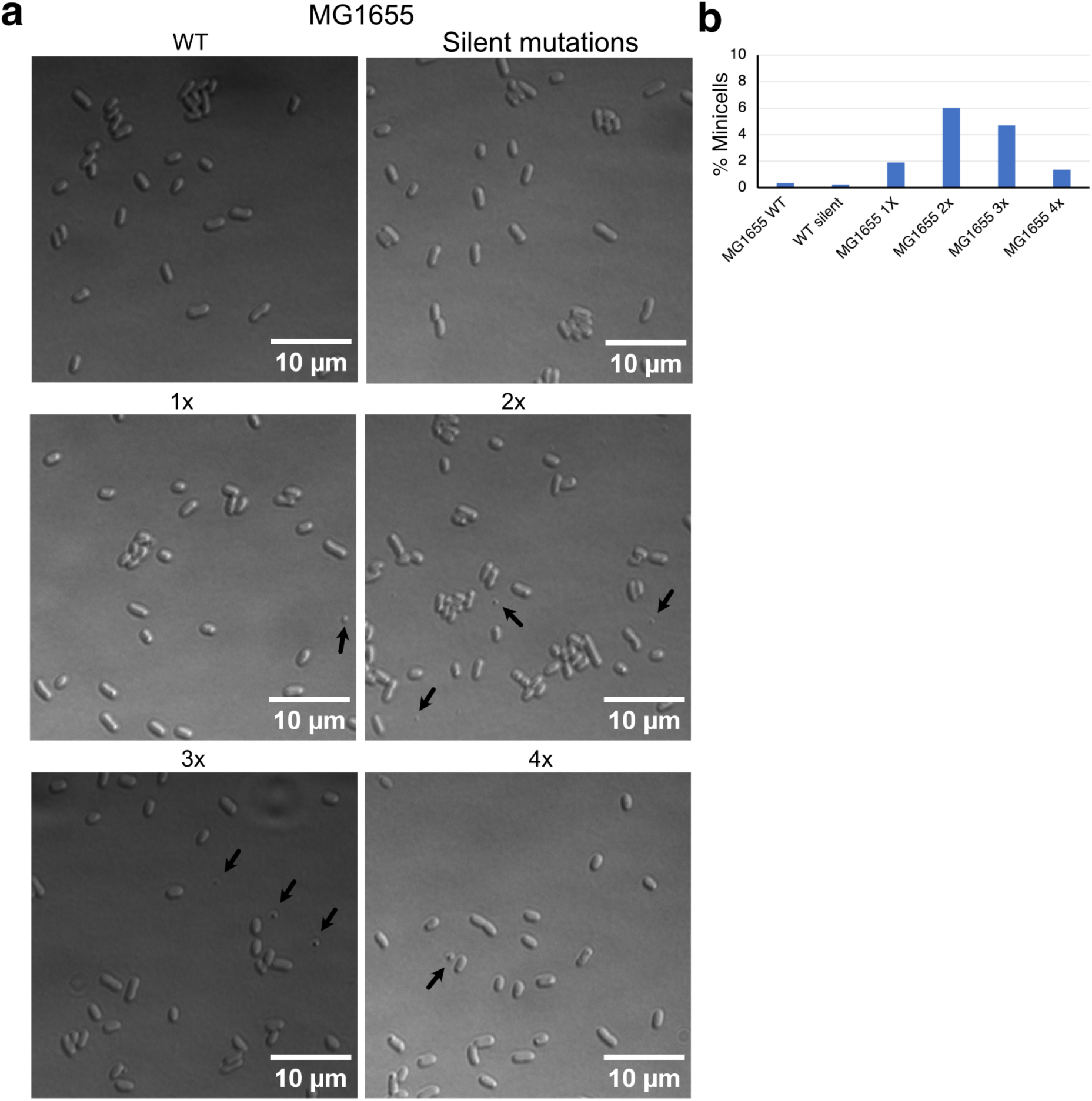
MinC linker mutants in the MG1655 background form few minicells. **a,** Representative DIC images of the indicated strains. Black arrows point to minicells (not all minicells are indicated). **b**, Bar graph showing the percentage of minicells among the total number of cells for the indicated strains.

**Fig. S8.**
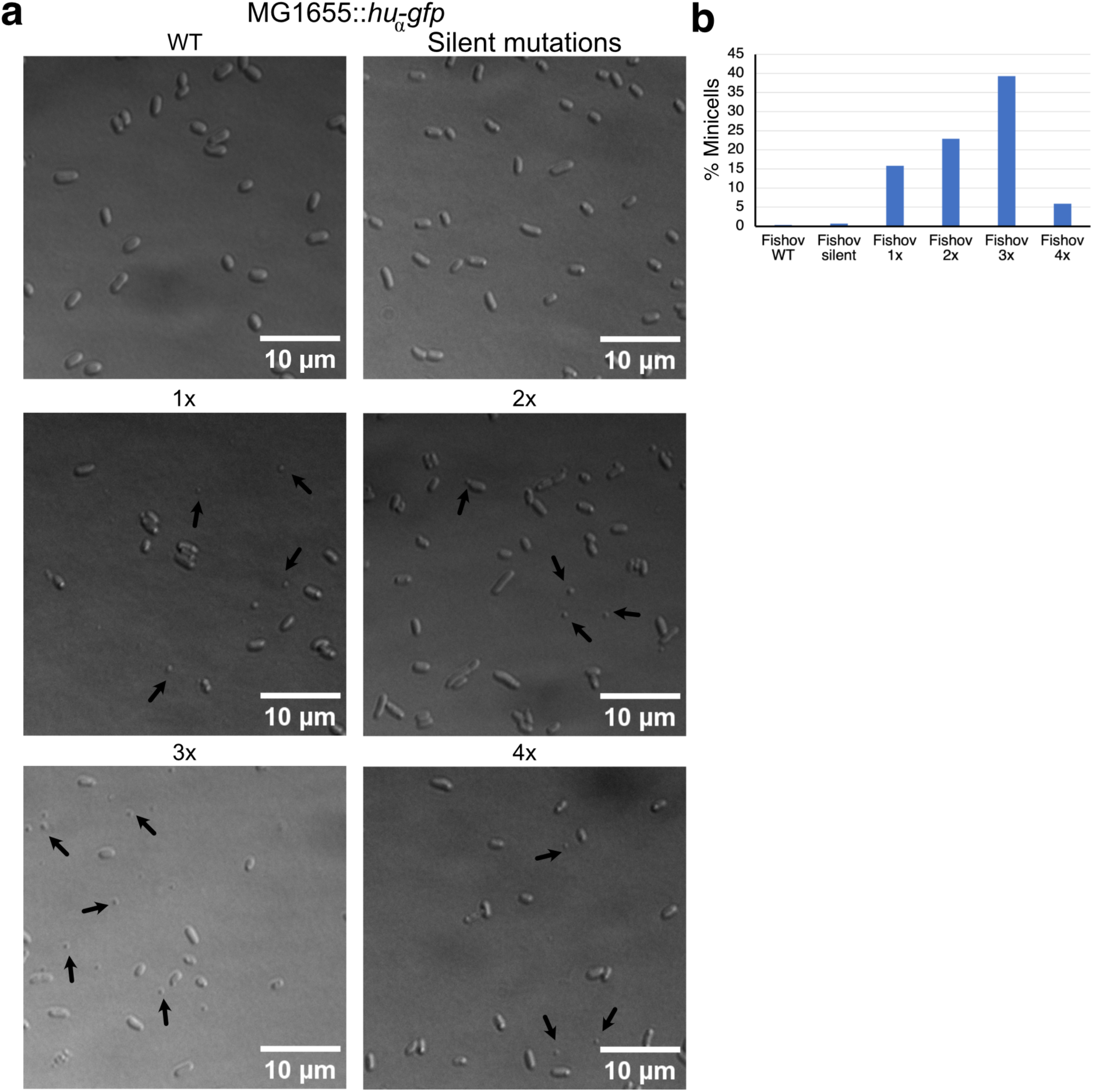
MinC linker mutants in the Fishov background form minicells. **a,** Representative DIC images of the indicated strains. Black arrows point to minicells (not all minicells are indicated). **b**, Bar graph showing the percentage of minicells among the total number of cells for the indicated strains.

**Fig. S9.**
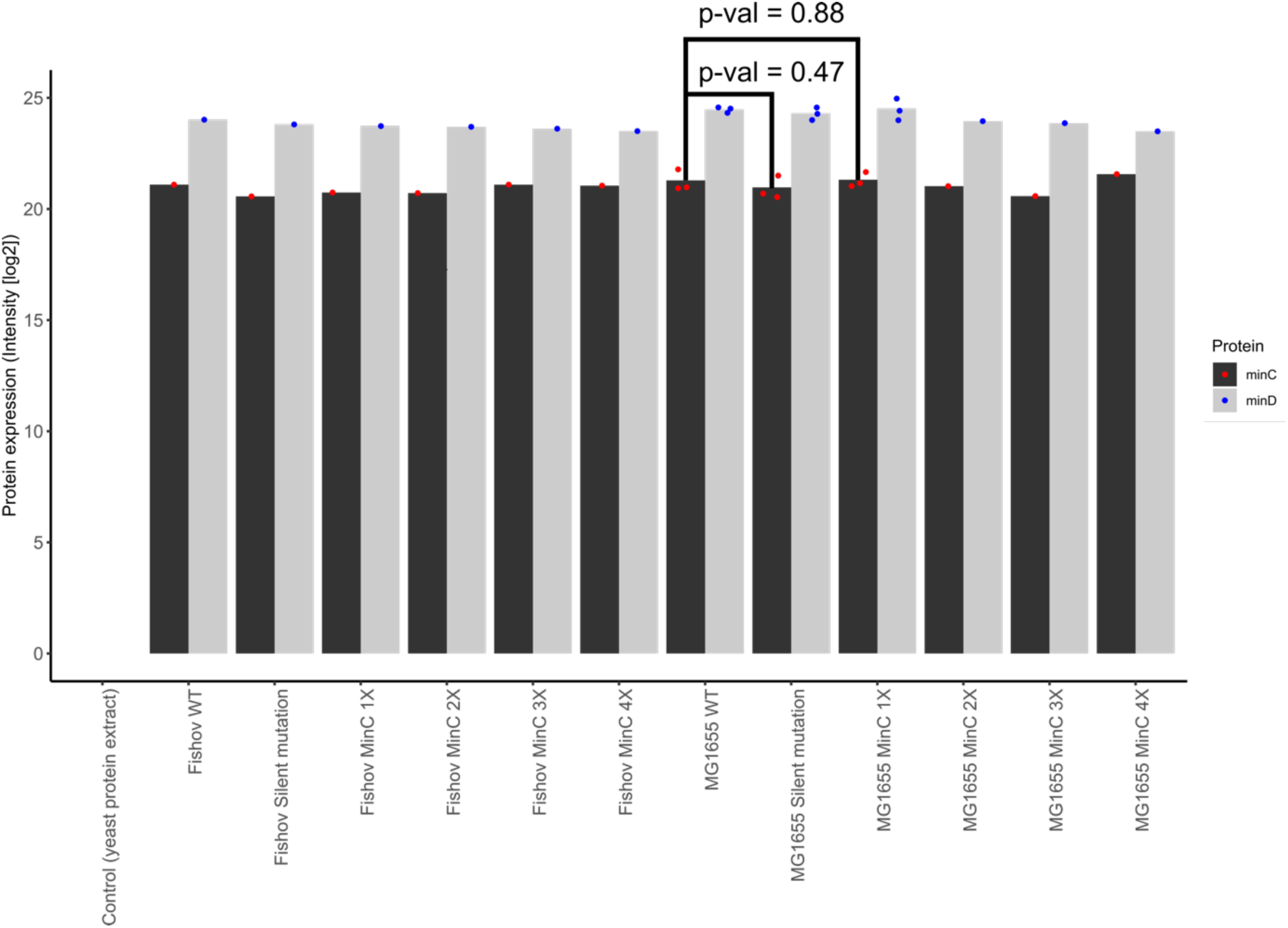
MinC linker mutants are expressed at similar levels as wild-type MinC. Bar graph showing the expression levels of MinC and MinD quantified by mass spectrometry. The x-axis indicates the names of the various strains, while the y-axis indicates the protein expression level, expressed as the log2 intensity of the proteins. Intensity refers to the signal strength corresponding to the relative abundance of a given protein in the sample. Each bar represents the summed intensity of a protein from two gel slices. For three constructs the experiment was repeated three times and the p-value was calculated. For more details, please see the methods section.

